# Macroevolutionary constraints on global microbial diversity

**DOI:** 10.1101/2022.06.04.494835

**Authors:** FJ Fishman, JT Lennon

## Abstract

Biologists have long sought to quantify the number of species on Earth. Often missing from these efforts is the contribution of microorganisms. Despite recent large-scale sampling efforts, estimates of global microbial diversity span many orders of magnitude. To reconcile this uncertainty, it is important to consider how speciation and extinction over the last four billion years constrain inventories of biodiversity. We parameterized macroevolutionary and mass-extinction event models to determine how diversification limits present-day microbial diversity. We find that while 10^6^-10^7^ taxa is most probable, much larger values (≥10^12^) are feasible. Allowing for mass extinction events does not greatly alter these conclusions. Along with empirical predictions, our models provide support for a massive global-scale microbiome while shedding light on the upper limits of life on Earth.

## INTRODUCTION

Global biodiversity is impossible to completely census, given the large number of individuals on the planet across a diverse range of habitats (Whitman *et al*. 1998). Various approaches have been used instead to approximate biodiversity, including diversity estimation based on partial censuses (Costello *et al*. 2012; Louca *et al*. 2019), ratios of taxonomic groupings (Mora *et al*. 2011), and many other macroecological and biogeographical methods (May 1988). The total number of species on the planet, when focusing on multicellular life, has been estimated to be ~10^6^-10^8^ species (Costello *et al*. 2012; Mora *et al*. 2011; Thompson *et al*. 2017). Despite their ubiquity, microorganisms have historically been overlooked in studies that attempt to estimate global biodiversity (Larsen *et al*. 2017). This oversight was largely due to technological reasons, as there were no comprehensive methods to systematically describe microbial diversity (Woese 1987).

With the advent of high-throughput DNA amplicon sequencing, large-scale quantification of microbial diversity became possible, yet the various approaches used to predict the total number of microbial species have generated highly divergent estimates (Larsen *et al*. 2017; Locey & Lennon 2016; Louca *et al*. 2019). Bacteria and archaea can be clustered into operational taxonomic units (OTUs) according to their similarity in the 16S rRNA gene, with the similarity cutoff often being set to 97% (Stackebrandt & Goebel 1994). While this clustering approach is often considered a conservative measure of bacterial and archaeal species (Eren *et al*. 2015; Poretsky *et al*. 2014), the 16S rRNA has served as a powerful tool, allowing microbiologists to survey the diversity of bacteria and archaea in a range of ecosystems across the planet (Thompson *et al*. 2017). Using the massive amount of data collected, several studies have attempted to quantify global bacterial and archaeal species diversity (*S*). One approach using collector’s curves estimated that microbes may only add 10^6^ species to the global estimates of plant and animal diversity (Louca *et al*. 2019), which means they would not fundamentally change current inventories, which are ~10^7^ at most (Mora *et al*. 2011). Another approach using the average number of unique bacterial species per host species estimated *S* to be approximately 10^9^ (Larsen *et al*. 2017; Wiens 2021). Last, a combination of scaling laws and biodiversity theory predicts that there are 10^12^ or more microbial taxa on Earth (Lennon & Locey 2020; Locey & Lennon 2016), including potentially 10^9^ in activated sludge systems alone (Wu *et al*. 2019). This ongoing debate is unresolved, as these predictions and estimates are difficult to directly test, but it may be possible to deem some values of present-day diversity as more likely than others.

These estimations and predictions should be considered in the context of constraints on present microbial diversity. The abundance of microorganisms at a global scale (*N*) has approached a steady state of 10^29^-10^30^ individuals (Kallmeyer *et al*. 2012; Whitman *et al*. 1998), and given that global taxon richness *S* cannot exceed the number of total individuals *N*, *S_present_* ≤ 10^30^ is a hard upper constraint on microbial richness. However, there may also be a soft upper constraint of 10^22^-10^23^ due to neutral drift, assuming (1) a constant 10^30^ bacterial and archaeal individuals and (2) a neutral drift rate of 4 – 5×10^-9^ substitutions per site per generation (Louca *et al*. 2019). A hard lower constraint of *S_present_* ≥ 10^6^ is also in place due to the number of reported 97% 16S rRNA OTUs (Schloss *et al*. 2016). Therefore, it is reasonable to surmise that the present number of bacterial and archaeal taxa *S_present_* is between 10^6^ and 10^23^.

Within this range, diversity is further constrained by macroevolutionary processes occurring over geological time scales. Speciation and extinction rates of lineages, the difference of which is the net diversification rate, should directly influence total present microbial diversity (Scholl & Wiens 2016) and determine the feasibility of both high and low estimates of this diversity. The simplest diversification models are birth-death processes, which assume constant and universal speciation and extinction rates (Raup 1985), but more complicated models should address realistic variation in these rates, such as clade-specific diversification rates (Moran *et al*. 1995; Scholl & Wiens 2016). In macro-organisms, well-documented mass extinction events are another such way the assumption of constant diversification is not upheld (Raup & Sepkoski 1982; Rohde & Muller 2005). These include the “Big Five” mass extinction events that eliminated 50-90% of marine invertebrate genera (Raup & Sepkoski 1982), as well as the Great Oxidation Event (GOE) (Gumsley *et al*. 2017), which likely caused the mass extinction of many lineages (Hodgskiss *et al*. 2019). Each of these mass extinction events may have reduced microbial diversity, thus constraining contemporary microbial richness, as a large portion of bacterial diversity is likely host-associated, (Hernández-Hernández *et al*. 2021; Thompson *et al*. 2017; Xie *et al*. 2005). The same factors causing the mass extinction of macro-organisms may also have elevated free-living microbial extinction (Newby *et al*. 2021), though the diversity of host-associated taxa should have been most greatly reduced. Models accounting for these phenomena may minimize uncertainty about the number of modern-day microbial taxa.

To understand how macroevolutionary rates constrain present-day species diversity, unbiased estimates of speciation and extinction are necessary. Any existing estimates of microbial diversification are derived from phylogenetic data (Louca *et al*. 2018; Scholl & Wiens 2016). Due to the nearly non-existent microbial fossil record, these phylogenies are constructed solely from molecular data, which may lead to incorrect rate estimation when diversification rates vary among lineages (Rabosky 2010; Stadler 2009). These phylogenies can also be generated by highly dissimilar birth-death processes that have divergent speciation and extinction dynamics (Louca & Pennell 2020). Such methods also require providing estimates for total microbial richness and the number of unsampled taxa to calculate diversification rates (Louca *et al*. 2018), which would run counter to the aim of using diversification rates to constrain present-day microbial richness. Therefore, diversification rate estimates that do not estimate unsampled taxa and that are not derived from molecular data alone are necessary to understand macroevolutionary constraints on species richness.

In this study, we seek to understand how speciation and extinction rates put additional constraints on present-day microbial diversity. To do so, we estimated speciation rates without phylogenetic inference to avoid the biases discussed above. With a simple model of diversification, we show the probability of various levels of present-day diversity. We then modify this model to account for mass extinction events to explore their potential effects on global diversity.

## METHODS

### Rate estimation

To consider bacterial and archaeal species, we must first consider our species definition. We phylogenetically defined a species as a cluster of strains with 97% 16S rRNA sequence similarity. While the 16S rRNA has limitations differentiating between certain species (Poretsky *et al*. 2014), its broad conservation across bacteria and archaea and relatively slow rate of evolution make it a convenient biomarker for considering global bacterial and archaeal diversity (Woese 1987). From this perspective, speciation occurs as the accumulation of 16S rRNA substitutions causes a focal sequence and an ancestral sequence to diverge by at least 3%. Thus, speciation rate *λ* can be calculated using 16S rRNA nucleotide substitution rates (*K*_16S_) as follows:

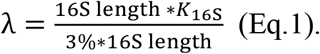

In Eq. 1, the numerator represents the total number of substitutions a 16S sequence undergoes over a million years, and the denominator represents the total number substitutions necessary for a 3% divergence in sequence, which is a speciation event by our OTU species definition.

To calculate λ values, we used a range of *K*_16S_ values (0.025–0.091% divergence/nt/My) based on the divergence of endosymbiotic bacteria in preserved and dated insects (Kuo & Ochman 2009). These *K*_16S_ values were calculated by calibrating the bacterial phylogenies with the age of their insect hosts, which possess a tractable fossil record (Moran *et al*. 1993). In this way, the ages of internal nodes of the bacterial phylogeny were mapped to corresponding ages in the insect phylogeny. These ages and the divergence between two bacterial 16S rRNA sequences were then used to directly calculate *K*_16S_. Using these *K*_16S_ values (Kuo & Ochman 2009), we calculated speciation rates of 0.0083–0.030 My^-1^. However, because substitution rates of endosymbiont bacteria are potentially twice that of their free-living relatives (Moran *et al*. 1995), we also considered speciation rates 50% smaller than the minimum endosymbiont-based speciation rate, producing a final range of 0.004–0.03 My^-1^. As an analogous technique cannot be used to estimate extinction rates (μ), we used values of relative extinction rates ε, the ratio of extinction to speciation (μ/λ), between 0 and 1 to account for various extinction scenarios.

### Expectations of birth-death process

The process of lineage diversification is often modeled as a stochastic birth-death process (Magallón & Sanderson 2001; Nee *et al*. 1994; Raup 1985), where speciation and extinction events are analogous to births and deaths of individuals, respectively. In diversification scenarios where present-day *S* ≥ 10^12^ taxa, the simulation of a stochastic birth-death process that stores times of birth and death events becomes computationally intractable. Due to this limitation, we first analyzed the expectations of birth-death processes *E*[*S_t_*] with constant speciation and extinction rates, which can be simply described by exponential growth when assuming the initial number of species to be 1:

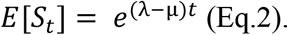

To compare the amount of diversity across various levels of λ and ε, we manipulated Eq. 2 into the following:

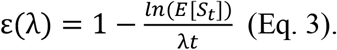

We plotted several contours of ε(*λ*) with various levels of *E*[*S_t_*] with *t* = 4000 My, a reasonable estimate of the time passed since the last universal common ancestor (LUCA) (Weiss *et al*. 2018). To calculate the probability of various ranges of present-day diversity, we calculated the area between contours via integration and normalized by the total area of feasible parameter space (10^6^ ≤ *E*[*S_t_*] < 10^23^).

### Mass extinction events

To understand the potential influence of mass extinction and its ability to constrain present-day microbial diversity, we considered the following mass extinction events: the Great Oxidation Event (GOE; ~2450 Mya) (Gumsley *et al*. 2017), the Ordovician-Silurian (O-S, 445 Mya); Devonian (D, 375 Mya); Permian-Triassic (P-Tr, 252 Mya); Triassic-Jurassic (Tr-J, 201 Mya); Cretaceous (K-T, 66 Mya) (Gumsley *et al*. 2017; Raup & Sepkoski 1982). Our model of mass extinction uses the expression for the expectations of a birth-death process (Eq. 2) and adds additional terms accounting for mass extinction events. We consider two new parameters influencing extinction beyond constant extinction rate μ: the intensity of mass extinction (*p*) and the proportion of taxa potentially affected by mass extinction (*q*). For each mass extinction event, there is a single reduction in the total number of species according to the magnitudes of *p* and *q*. To obtain the number of species at time *t*, we multiply the birth-death expectations (Eq. 2) by the proportion of taxa surviving each mass extinction event (1 – *pq*) for as many mass extinction events occurring by time *t*, where *t* is some nonnegative integer:

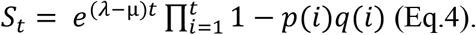

Let *M* be the set of the six timesteps where mass extinction occurred, and *M_H_* be the set of timesteps with host-associated mass extinction. Mass extinction intensity *p*(*i*) is equal to some value *p* during a mass extinction event and 0 for all other timesteps:

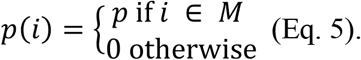

We consider situations where *p* is 0.0, 0.5, or 0.9 to model situations without mass extinction, with moderate mass extinction, and with intense mass extinction, respectively. Likewise, the proportion of taxa vulnerable to mass extinction *q*(*i*) at timestep *i* is set to some value *q* during a host-associated mass extinction event and 1 for all other timesteps:

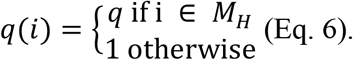

Therefore, the proportion of species removed due to mass extinction is *pq* during host-associated mass extinction, *p* during non-host-associated mass extinction, and 0 for all other timesteps. When considering Eqs. 5 and 6 when *t* = 4000 My (present day), the Great Oxidization Event is the only non-host-associated mass extinction event, so there is one timestep where the proportion of species removed is *p* and five timesteps where the proportion is *pq*. Eq. 4 then becomes

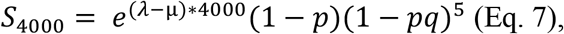

which we further transformed to produce contours of *ε* in terms of *λ:*

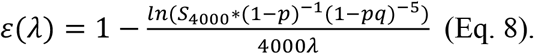

To obtain informed estimates of *q*, we assumed that host-associated bacteria and archaea would be more likely to go extinct from macro-organismal mass extinction than would free-living microbes. In place of the unmeasurable ancient proportion of host-associated species, we calculated the present-day proportion of host-associated microbial species, as well as obligately host-associated and preferentially host-associated proportions, using species and site data from the Earth Microbiome Project (EMP) (Thompson *et al*. 2017). We defined obligately host-associated taxa as OTUs only sampled from hosts and preferentially host-associated taxa as those found in hosts for over 50% of their total occurrences.

## RESULTS

### Expectations of birth-death process

To evaluate how diversification parameters constrain present-day microbial species richness, we expressed relative extinction rate (ε) as a function of speciation rate (λ) at various contours of *E*[*S_t_*] (Eq. 3) and with *t* = 4000 My (present-day) within the bounds of the speciation rate range described above (Fig. 1). Expected diversity increases as extinction decreases and speciation rise, though the relationship between ε and λ is non-linear (Eq. 5). Approximately 50% of the combinations of *λ* and *ε* lead to infeasibly low (<10^6^) or high (>10^23^) diversity (Table S1). Certain high levels of diversity require *ε* to be sufficiently low and λ to be sufficiently large (Fig. 1A, B). For instance, 10^12^ species are only possible for ε < 0.78 and λ > 0.007 sp./My. However, there are much less strict limitations on ε or λ for reaching 10^6^ species. In terms of amount of feasible parameter space, lower diversity outcomes are somewhat more likely than high diversity outcomes (Fig. 1C). For instance, the probability of 10^6^-10^7^ species is ~9.0%, while the probability of 10^12^-10^13^ species is ~6.2%. A reason for the decrease in probability for higher levels of diversity is that it is impossible to reach them at relatively low values of λ (Fig. 1B). However, once λ reaches ~0.013 sp./My, any further increase in λ does not alter the relative probabilities of each outcome (Fig. 1B). However, each of these ranges of diversity is well within these constraints and far from extreme outcomes, and no outcome is far more probable than the others. With this simple analysis, we show that vast diversity is indeed possible within the specified speciation constraints.

**Fig. 1.**
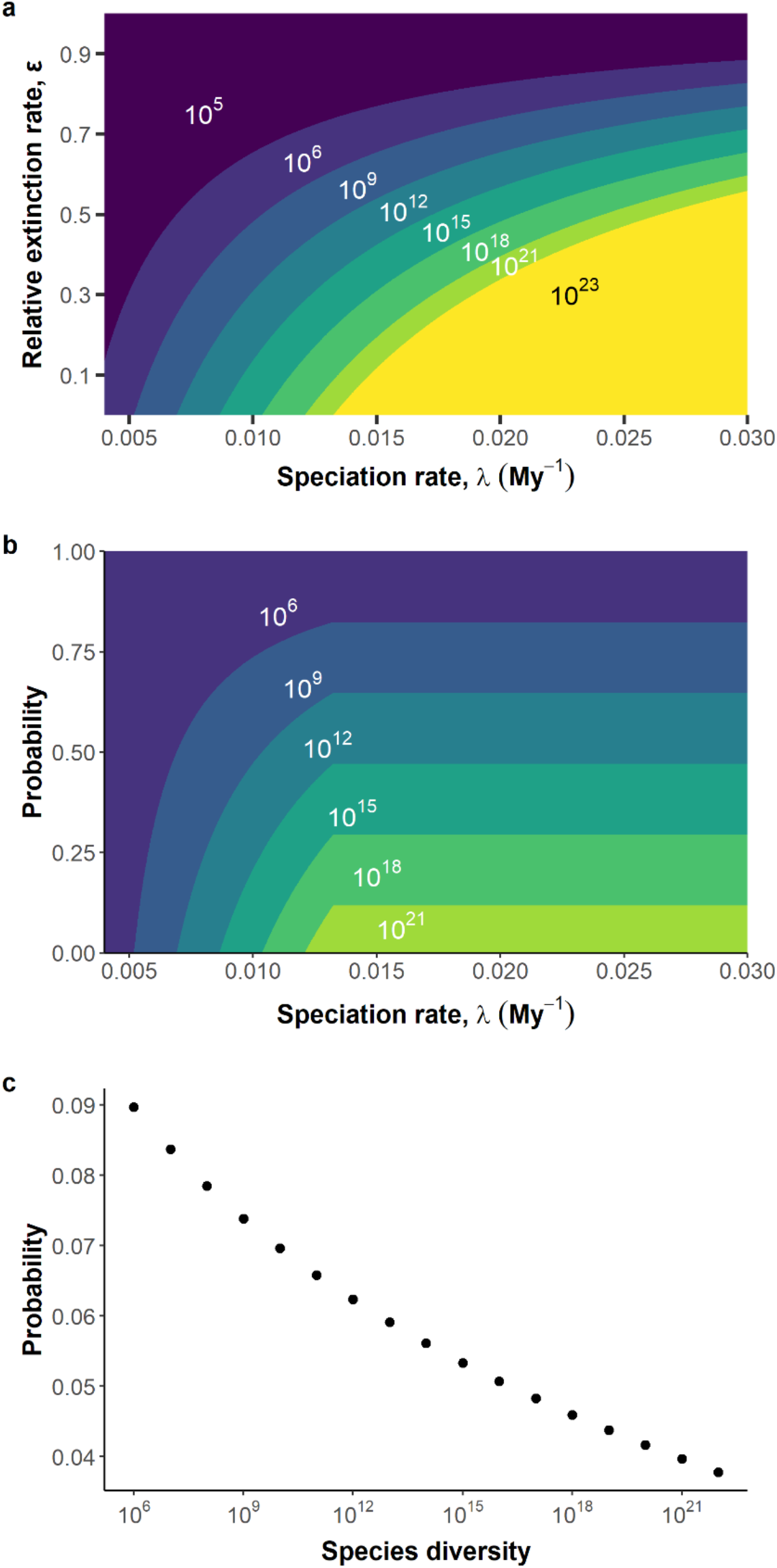
Expected present-day diversity (*E*[*S*_4000_]) of a birth-death process and probabilities of certain outcomes. (A) Combinations of speciation (0.004 ≤λ≤ 0.03) and relative extinction rates (0 ≤ *ε* ≤ 1) lead to a wide range of *E*[*S*_4000_]. The regions labeled 10^5^ and 10^23^ consist of combinations of diversification parameters that lead to infeasibly low (*S* < 10^6^) and infeasibly high (*S* > 10^23^) species diversity, respectively. All other labeled regions correspond to *E*[*S*_4000_] within a three order of magnitude bin (e.g., 10^6^ label corresponds to 10^6^-10^9^ species) except 10^21^ (10^21^-10^23^ species). Each labeled region is separated by contours of ε(λ) (Eq. 3) with values of *E*[*S*_4000_] between 10^6^ and 10^23^ species. (B) The probability of diversity outcome bins across speciation rate. The probability is calculated as the proportion of *ε* values leading to a diversity outcome (e.g., 10^6^-10^9^ species) at a given value of *λ*. (C) The overall probability of diversity outcomes spanning one order of magnitude. This probability was calculated by creating contours of ε(λ) (Eq. 3) with *E*[*S*_4000_] set from 10^6^ to 10^23^ and calculating the area between each contour and normalizing by the total area of the feasible parameter space.

### Mass extinction events

Our simulations show that the intensity of mass extinction events determines their effect on present microbial biodiversity (Fig. 2). Let us first consider scenarios where mass extinction affects all microbes equally (*q* = 1) and strongly (*p* = 0.9; Fig. 2A, B). Compared to scenarios without mass extinction, much higher speciation rates and lower extinction rates are required to reach equivalent levels of species diversity (Fig. 1A, 2B). For example, net diversification parameters resulting in 10^12^ species when *p* = 0 lead to ~10^6^ species when *p* = 0.9 (Fig. 2A). Additionally, the proportion of total feasible parameter space decreases from 50.6% in the birthdeath expectation model to 36.5% in the mass extinction model (Table S1). The proportion of parameter space leading to >10^6^ species increases to 50.2% from 26.7%, as well (Table S1). However, the relative probabilities of each diversity outcome within the feasible parameter space are primarily unchanged (Fig. S1). Setting *p* = 0.9 is comparable to the degree of extinction in macro-organisms during the Permian-Triassic, the most severe mass extinction event (Sepkoski 1990). If these events lead to even a 50% diversity reduction, present-day diversity still is decreased, though the effect is diminished compared to the 90% scenario. However, it is clear that severe mass extinction can greatly change the outcomes for individual parameter combinations.

**Fig. 2.**
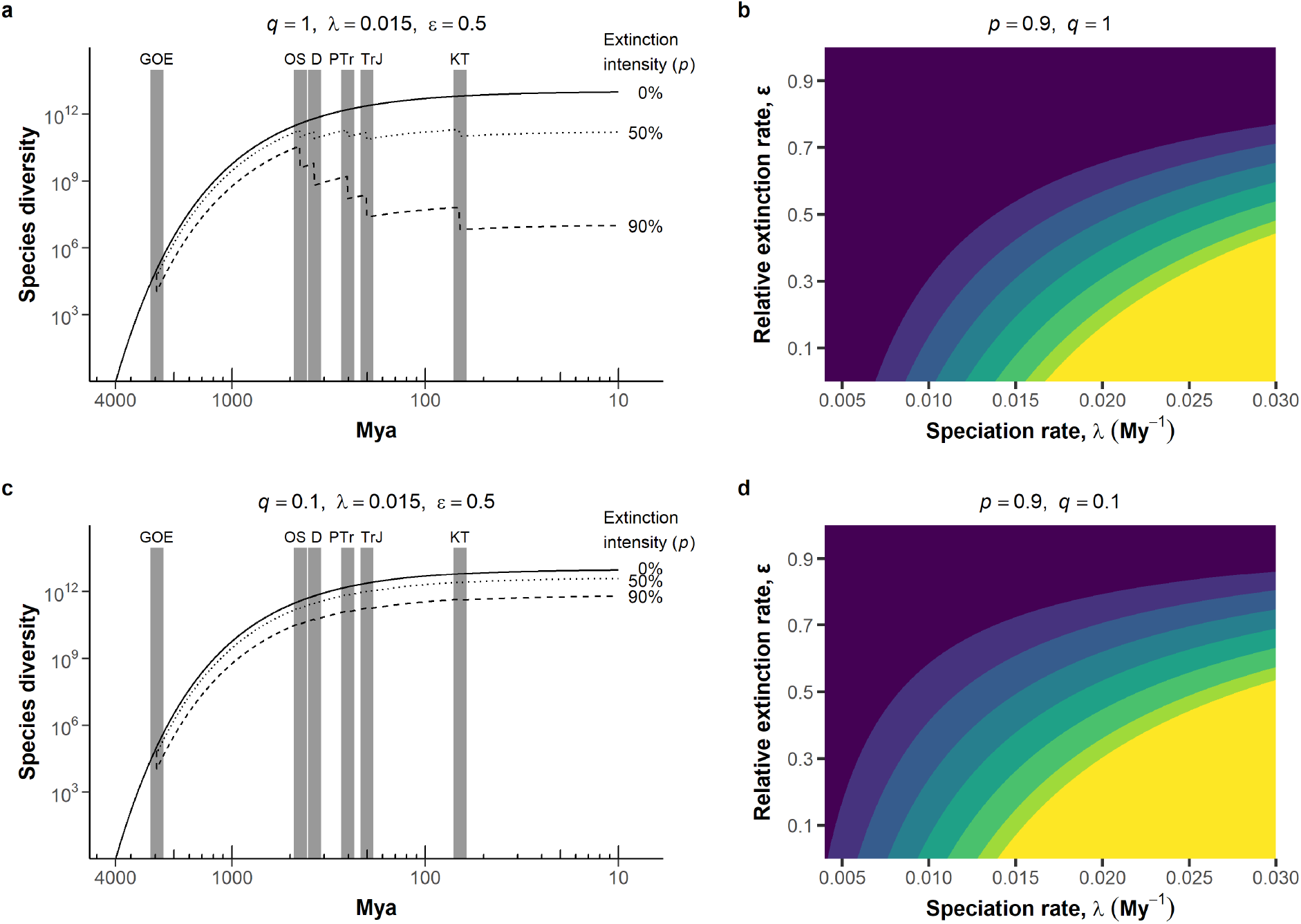
Effect of mass extinction events on species diversity (A, B) when all taxa are vulnerable to mass extinction (*q* = 1) and (C, D) when only obligate host-associated taxa are vulnerable (*q =* 0.1). (A, C) Species diversity over time was calculated using the mass extinction model (Eq. 4) with speciation and relative extinction at λ = 0.015 My^-1^ and *ε* = 0.5, respectively. Known mass extinction events (grey bars) cause a one-time 0% (solid), 50% (dotted), or 90% (dashed) reduction in diversity of vulnerable species at each mass extinction event. (GOE: Great Oxidation Event; OS: Ordovician-Silurian; D: Devonian; PTr: Permian-Triassic; TrJ: Triassic-Jurassic; KT: Cretaceous). (B, D) Present-day diversity calculated from mass extinction model across a range of speciation (0.004 ≤ λ ≤ 0.03) and relative extinction rates (0 ≤ *ε* ≤ 1) with mass extinction intensity *p* = 0.9. Each labeled region (same coloration scheme as Fig. 1A) is separated by contours of ε(λ) (Eq. 8) with values of *E*[*S_present_*] between 10^6^ and 10^23^.

To obtain an informed estimate of the proportion of taxa vulnerable to mass extinction (*q*), we used species and site data from the Earth Microbiome Project (EMP) to calculate the present-day proportion of host-associated 16S OTUs. We found that ~90% of EMP bacterial OTUs were free-living to some degree, meaning that only ~10% of EMP OTUs were obligately host-associated (Table 2). However, about half of all OTUs were host-associated to some degree. Therefore, if mass extinction events of macro-organisms only resulted in the extinction of host-associated organisms, only between 10% and 50% of microbial taxa would be vulnerable to mass extinction, depending on if all host-associated species, only obligately host-associated species, or some mixture is considered.

**Table 1.** Defining key model parameters used in birth-death process and mass extinction models.

| Variable/Parameter | Description |
| --- | --- |
| Global species diversity ($S_t$) | The total number of 97% 16S rRNA bacterial and archaeal OTUs present on the planet at time $t$. |
| Speciation rate ($\lambda$) | The number of species an extant species generates per million years ($\text{My}^{-1}$). |
| Extinction rate ($\mu$) | Number of species extinctions per extant species per million years ($\text{My}^{-1}$). |
| Relative extinction rate ($\epsilon$) | Ratio of extinction rate to speciation rate ($\mu / \lambda$). |
| Mass extinction intensity ($p$) | The proportion of vulnerable species removed at a certain timestep from a mass extinction event. |
| Vulnerable proportion of taxa ($q$) | The proportion of species that are vulnerable to mass extinction. |
| Mass extinction events ($M$) | The set of timesteps where mass extinction occurs. |
| Host-associated mass extinction events ($M_H$) | The set of timesteps where mass extinction of hosts occurs. |

**Table 2.** The proportion of Earth Microbiome Project (EMP) 16S rRNA OTUs classified as host-associated or free-living. Obligate host-associated OTUs are only found in samples taken from hosts, as opposed to preferentially host-associated (over 50% of samples from hosts) and host-associated OTUs (any samples are from hosts). The same terms are applied to free-living OTUs.

| Niche | Proportion of all EMP OTUs |
| --- | --- |
| Host-associated | 47.8% |
| Preferentially host-associated | 19.9% |
| Obligate host-associated | 9.3% |
| Free-living | 90.7% |
| Preferentially free-living | 78.9% |
| Obligate free-living | 52.2% |

When we consider scenarios when only a fraction of microbial species is affected by mass extinction, the model begins to converge to the expectations of the birth-death processes (Fig. 2C, D). With *q* = 0.1, which corresponds to only 10% of microbial lineages being vulnerable to mass extinction, mass extinction has a much more muted effect. The same net diversification parameters described above resulting in 10^12^ species when *p* = 0 now leads to >10^10^ species at *p* = 0.9. The feasible parameter space also converges to that of the birth-death expectations as well (Table S1, Fig. 1A, 2D). We also modeled scenarios where only certain groupings of lineages were vulnerable to mass extinction and identified scenarios where this allows for present-day diversity on the order of the birth-death expectations (Fig. S2), illustrating another scenario where adding biological detail can reduce the effect of mass extinction. Therefore, while extreme mass extinction scenarios may constrain microbial diversity, more conservative scenarios suggest that mass extinction may have only moderately decreased present-day diversity.

## DISCUSSION

In this study, we modeled how macroevolutionary rates influence present-day microbial species diversity in an attempt to further constrain the estimates and predictions from previous studies ranging from 10^6^ to 10^12^ species (Larsen *et al*. 2017; Locey & Lennon 2016; Louca *et al*. 2019; Wiens 2021). The values suggested in these studies all can be generated from feasible combinations of macroevolutionary rates (Fig. 1,4). In fact, our results introduce the possibility that bacterial and archaeal diversity may outstrip the largest predictions (Lennon & Locey 2020; Locey & Lennon 2016). Given the diversification parameters and the model we used, 10^6^ species is more likely than 10^9^, both of which are more likely than 10^12^ (Fig. 2,3). However, while our study finds that it is most likely that total present-day diversity is not orders of magnitude larger than current inventories, it does not deem any previously made prediction or estimate vanishingly unlikely or even improbable (Larsen *et al*. 2017; Lennon & Locey 2020; Locey & Lennon 2016; Louca *et al*. 2019; Wiens 2021).

The simple approach we use in this study is not without its caveats and assumptions. This study only uses substitution rate data from obligately host associated taxa, which may not be representative of the overall rate of molecular evolution of free-living microbes (Espejo & Plaza 2018; Moran *et al*. 1995). While a more representative sample of substitution rates from free-living lineages or taxa with multiple 16S rRNA copies may improve upon this study, such data are scarce. Thus, our analysis provides a basis for feasible levels of microbial diversity with backing from the fossil record and including speciation values that account for the differences between free-living and obligate endosymbiont substitution rates (Moran *et al*. 1995). Additionally, we did not attempt to directly estimate extinction rates, as microbial extinction cannot reasonably be estimated apart from phylogenetic approaches relying on *a priori* assumptions of microbial richness (Louca *et al*. 2018), which given the objectives of our study would introduce circular reasoning. If an unbiased method for estimating relative extinction was found, then it could be used to further constrain the diversification parameter space. Our analysis also assumes no biogeographical or niche association with diversification (Li & Wiens 2022). It is quite likely that clades associated with certain environments have higher diversification rates than those of other environments (Rabosky 2020). While clade-specific diversification rates were modeled here, a more thorough modeling process including diversification dynamics of specific bacterial lineages may provide more insight into global diversification.

Our simulations of mass extinction events showed that while severe mass extinction can constrain present-day diversity, there are many scenarios that result in little change compared to our model without mass extinction. This convergence to birth-death expectations occurs as the proportion of lineages affected by mass extinction decreases. In fact, the true proportion of bacteria affected by host mass extinction may have been smaller than the proportion of obligately host-associated taxa depending on the host range of the microbial lineages. For instance, if one microbial taxon is present in several host taxa, extinction is unlikely if only one host taxon becomes extinct. However, the publicly available 16S rRNA databases do not typically contain information regarding whether OTUs were found in a narrow or broad range of host taxa, only the general source of each sample. It is also possible for plant and animal mass extinction to affect more than just host-associated microbes if higher-order effects of extinction of macro-organisms had downstream effects on free-living microbes, thus increasing the possible percentage of microbial taxa vulnerable to mass extinction. However, explicitly modeling such effects here is unnecessary, as the outcomes with high *q* will simply converge to our first mass extinction scenario with *q* = 1.0.

Our mass extinction model contains other assumptions and caveats as well. To simplify the model, we implemented mass extinction as a one-time reduction in diversity per event. These events might be more realistically modeled as occurring over the span of several million years. We implemented each of the “Big Five” mass extinction events as equal in extinction magnitude, but some of these events had larger effects on host diversity than others (Raup & Sepkoski 1982), which likely would have scaled onto microbial extinction. Additionally, there may have been other microbe-specific mass extinction events besides the GOE that could have had a profound impact on diversity. Our models also do not take into consideration increases in diversification via adaptive radiation following mass extinction (Stroud & Losos 2016). Despite these caveats, our models provide a foundation for how losing large proportions of diversity several times may have altered present-day diversity by examining extreme scenarios.

Our study finds vast diversity beyond 10^12^ species is indeed possible and only marginally less likely than lower levels of diversity. While this analysis suggests the globe is most likely to contain fewer than 10^8^ microbial species, our approach cannot make a precise prediction on microbial diversity, nor can it rule out the predictions and estimates made by previous studies (Larsen *et al*. 2017; Lennon & Locey 2020; Locey & Lennon 2016; Louca *et al*. 2019; Wiens 2021). The simple models described here use speciation rates calculated from endosymbiotic bacterial 16S rRNA substitution rates, which do not have the inherent bias of requiring estimates of unsampled taxa. These models provide a new angle with which to address the question of global microbial diversity. New approaches will be necessary to confront the lack of consensus in the field as we seek to reconcile the estimations and results put forth, such as methods going beyond 16S rRNA-based species definitions and embracing the ecological and functional differences among micro-organisms (Arevalo *et al*. 2019). Such approaches may reveal levels of diversity greater than any suggested by 16S rRNA approaches.

## ACKNOWLEDGEMENTS

An earlier version of the manuscript was improved based on critical feedback from members of the Lennon lab, including discussions with KJ Locey. Research was supported by the National Science Foundation (DEB-1934554 JTL, DBI-2022049 JTL), US Army Research Office Grant (W911NF-14-1-0411 JTL), the National Aeronautics and Space Administration (80NSSC20K0618 JTL). Data and code can be found here: https://github.com/LennonLab/MicroSpeciation.

## Notes

https://github.com/LennonLab/MicroSpeciation

### Competing Interest Statement

The authors have declared no competing interest.

