## Supplement for "Macroevolutionary constraints on global microbial diversity"

Fishman FJ, Lennon JT*

*Unequal diversification among lineages*

To simulate clade-specific diversification, we implemented a model allowing for the existence of clades with different sensitivities to mass extinction events. We initiate the model with a single clade containing one species. This clade experiences $p_0$ = 90% loss in diversity due to mass extinction as well as background extinction $\mu$, as in our mass extinction model. With probability $c$, a new clade $i$ with a single species arises from the initial clade with the same relative extinction rate, but $p_i$, the proportion of diversity this clade loses due to mass extinction, is drawn from a uniform distribution from 0 to 1. Both of these clades can now generate new clades with probability $c$ per clade, causing the rate of clade generation per timestep to increase with the number of clades present. This leads to the generation of many clades with various degrees of sensitivity to mass extinction events.

Table S1. The percentage of parameter space ($0.004 \leq \lambda \leq 0.03$, $0 \leq \varepsilon \leq 1$) leading to infeasibly low ($S < 10^6$), feasible ($10^6 \leq S \leq 10^{23}$), and infeasibly high levels ($S > 10^{23}$) of present-day species diversity for the expectations of a birth-death process and the mass extinction model ($p = 0.9$) with all species vulnerable to mass extinction ($q = 1$) and with only obligately host-associated species vulnerable ($q = 0.1$).

| Model | Infeasibly low diversity $S < 10^6$ | Feasible diversity $10^6 \leq S \leq 10^{23}$ | Infeasible high diversity $S > 10^{23}$ |
|---|---|---|---|
| Birth-death expectations | 26.7% | 50.6% | 22.8% |
| Mass extinction ($q = 1$) | 50.2% | 36.5% | 13.5% |
| Mass extinction ($q = 0.1$) | 32.1% | 47.2% | 20.7% |

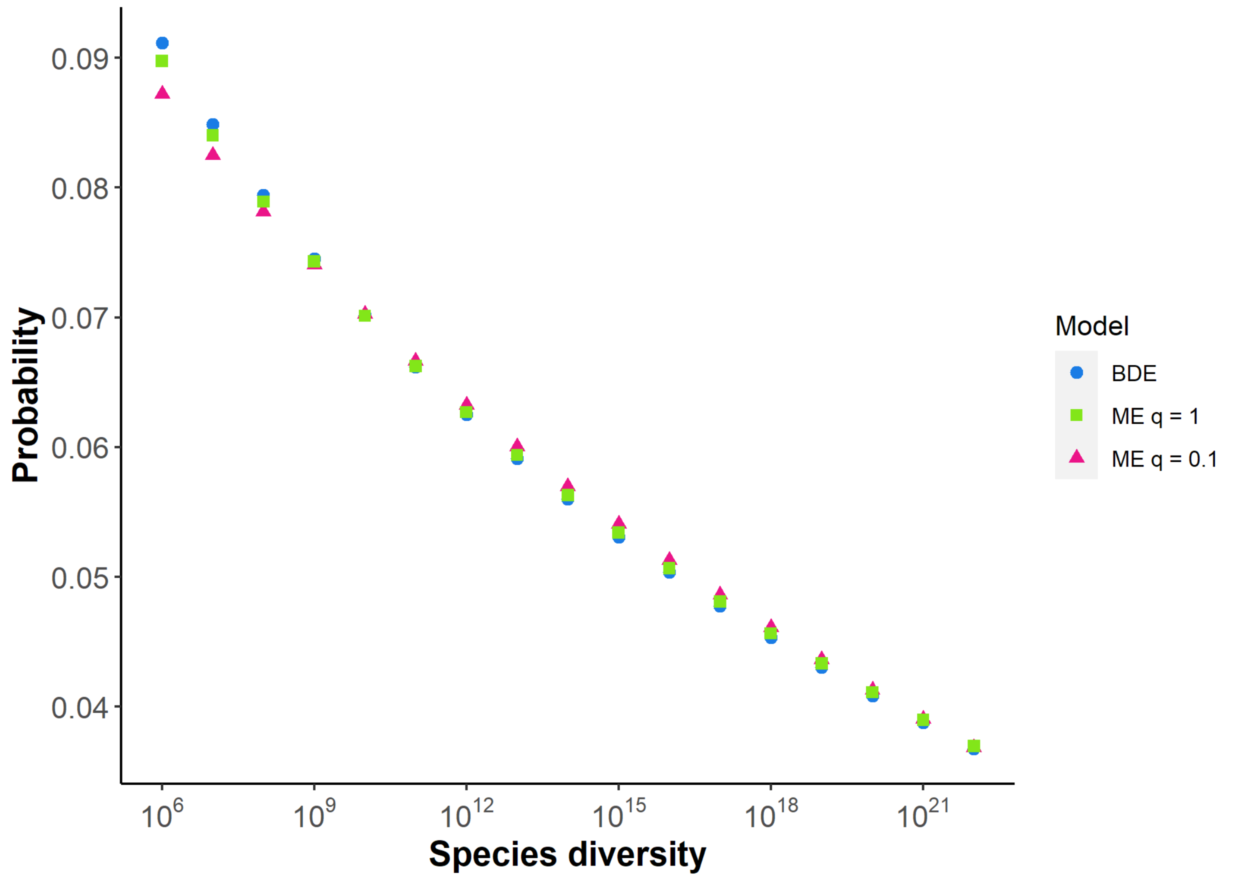


Fig. S1. The overall probability of diversity outcomes spanning one order of magnitude for the birth-death expectations model, the mass extinction model with all species vulnerable ($q = 1$), and the mass extinction model with only obligately host-associated species vulnerable. These probabilities were calculated by creating contours of $\varepsilon(\lambda)$ (Eq. 8) with $S_{4000}$ set from $10^6$ to $10^{23}$ and calculating the area between each contour and normalizing by the total area of the feasible parameter space for each model

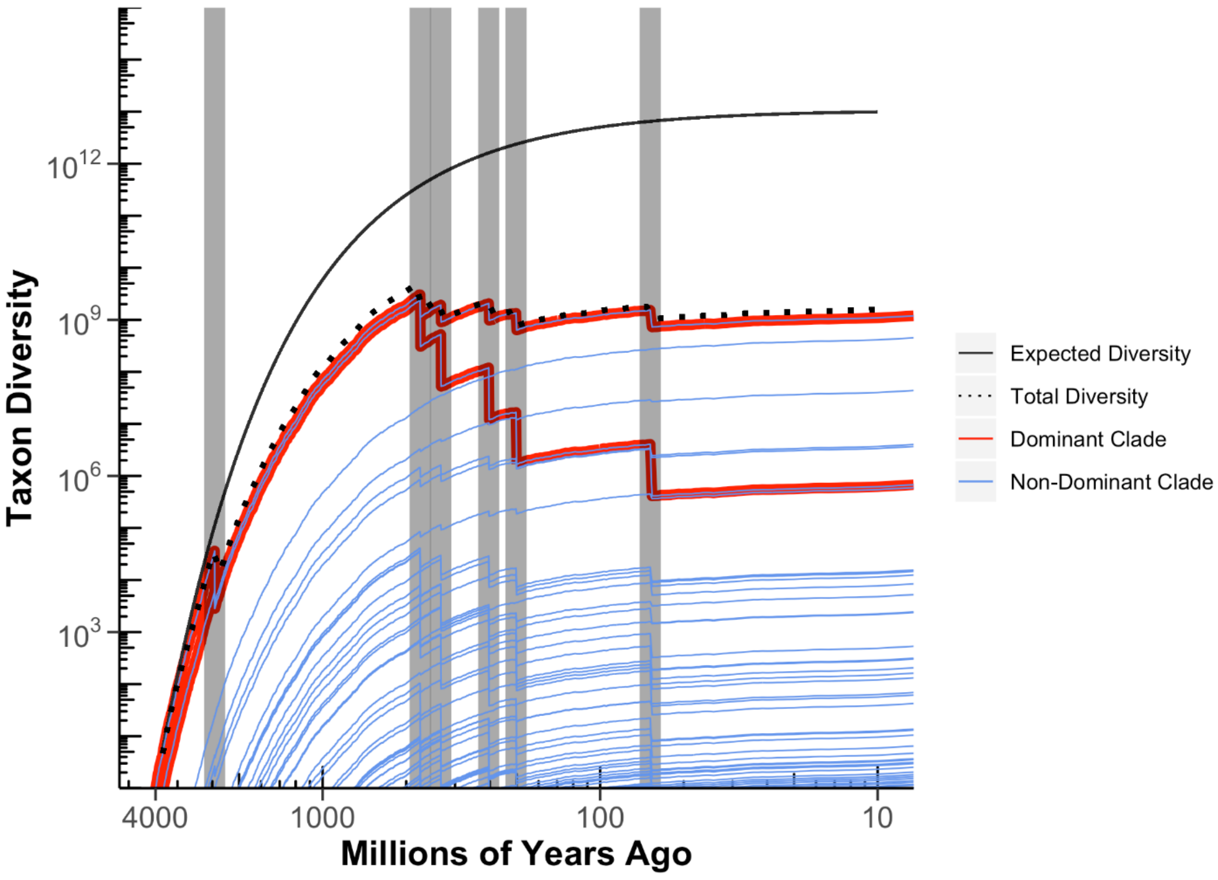


Fig. S2. The effect of clade-specific diversification on present-day species diversity. Each clade is displayed as a blue curve, and the total diversity is displayed as a dotted line. Dominant clades that at one point have the highest richness are highlighted in red. The solid black line represents the expected diversity of birth-death process with the same diversification parameters ($\lambda = 0.015$, $\varepsilon = 0.5$. Here, while the initial dominant clade is very strongly affected by mass extinction (~$10^6$ present-day species), another clade that becomes dominant is less strongly affected (~$10^9$ present-day species) despite the precipitous decline of the original clade. Unequal diversification among lineages here results in high total microbial diversity to be maintained in the system.
